# Genetic toolkit for sociality predicts castes across the spectrum of social complexity in wasps

**DOI:** 10.1101/2020.12.08.407056

**Authors:** Christopher D. R. Wyatt, Michael Bentley, Daisy Taylor, Ryan E. Brock, Benjamin A. Taylor, Emily Bell, Ellouise Leadbeater, Seirian Sumner

**Author notes:** Corresponding authors, **Christopher Douglas Robert Wyatt**, Centre for Biodiversity and Environment Research, University College London, London, UK., **Seirian Sumner**, Centre for Biodiversity and Environment Research, University College London, London, UK. Equal contribution (Michael Bentley, Daisy Taylor).

## Abstract

Major evolutionary transitions describe how biological complexity arises; e.g. in evolution of complex multicellular bodies, and superorganismal insect societies. Such transitions involve the evolution of division of labour, e.g. as queen and worker castes in insect societies. Castes across different evolutionary lineages are thought to be regulated by a conserved genetic toolkit. However, this hypothesis has not been tested thoroughly across the complexity spectrum of the major transition. Here we reveal, using machine learning analyses of brain transcription, evidence of a shared genetic toolkit across the spectrum of social complexity in Vespid wasps. Whilst molecular processes underpinning the simpler societies (which likely represent the origins of social living) are conserved throughout the major transition, additional processes appear to come into play in more complex societies. Such fundamental shifts in regulatory processes with complexity may typify other major evolutionary transitions, such as the evolution of multicellularity.

## Main

The major evolutionary transitions span all levels of biological organisation, facilitating the evolution of life’s complexity on earth via cooperation between single entities (e.g. genes in a genome, cells in a multicellular body, insects in a colony), generating fitness benefits beyond those attainable by a comparable number of isolated individuals^1^. The evolution of sociality is one of the major transitions and is of general relevance across many levels of biological organisation from genes assembled into genomes, single-cells into multi-cellular entities, and insects cooperating in superorganismal societies. The best-studied examples of sociality are in the hymenopteran insects (bees, wasps and ants) - a group of over 17,000 species, exhibiting levels of sociality across the transition from simple sociality (with small societies where all group members are able to reproduce and switch roles in response to opportunity), through to complex societies (consisting of thousands of individuals, each committed during development to a specific cooperative role and working for a shared reproductive outcome within the higher-level ‘individual’ of the colony, known as the ‘superorganism’^2^). Recent analyses of the molecular mechanisms of insect sociality have revealed how conserved suites of genes, networks and functions are shared among independent evolutionary events of insect superorganismality^3–7^. An outstanding question is to what extent are genomic mechanisms operating *across levels* of complexity in the major transition – from simple to complex sociality – conserved^8^? A lack of data from representatives across any one lineage of the major transition have limited our ability to address this question.

A key step in the evolution of sociality is the emergence of a reproductive division of labour, where some individuals commit to reproductive or non-reproductive roles, known as queens and workers respectively in the case of insect societies. An overarching mechanistic hypothesis for social evolution is that the repertoire of behaviours typically exhibited in the life cycle of the solitary ancestor were uncoupled to produce a division of labour among group members with individuals specialising in either the reproductive (‘queen’) or provisioning (‘worker’) phases of the solitary ancestor^9^. Such phenotypic decoupling implies that there will be a conserved mechanistic toolkit that regulates queen and worker phenotypes in species representing different levels of social complexity across the spectrum of the major transition (reviewed in^10^). An alternative to the shared toolkit hypothesis is that the molecular processes regulating social behaviours in non-superorganismal societies (where caste remains flexible, and selection acts primarily on individuals) differ fundamentally from those processes that regulate social behaviours in superorganismal societies ^11,12^. Phenotypic innovations across the animal kingdom have been linked to genomic evolution: taxonomically-restricted genes^13–16^, rapid evolution of proteins^17,18^ and regulatory elements^17,19^ been found in most lineages of social insects^20^. Indeed, some recent studies have suggested that the processes regulating different levels of social complexity may be different^17,19,21^. The innovations in social complexity and the shift in the unit of selection (from individual-to group-level^22^) that accompany the major transition may therefore be accompanied by genomic evolution, throwing into question whether a universal conserved genomic toolkit regulates social behaviours across the spectrum of the major transition^8^. The roles of conserved and novel processes are not necessarily mutually exclusive; novel processes may coincide with phenotypic innovations, whilst conserved mechanisms may regulate core processes at all stages of social evolution.

Currently, data are largely limited to species that represent either the most complex – superorganismal - levels of sociality (e.g. ants or honeybees^23^), or the simplest levels of social complexity as non-superorganisms that likely represent the first stages in the major transition (e.g. *Polistes* wasps^7,24–26^ and incipiently social bees^27–30^). We lack data on the intermediary stages of the major transition and thus lack a comprehensive analysis of if and how molecular mechanism change *across* any single evolutionary transition to superorganismality. One exception is a recent study that identified a core gene set that consistently underlie caste-differentiated brain gene expression across five species of ants^5^; however, this study lacked ancestrally non-superorganismal representatives (one species had secondarily lost the queen caste but evolved from a superorganismal ancestor^7^).

A promising group for exploring these questions are the social wasps^31^, with some 1,100 species exhibiting the full spectrum of sociality. We generated brain transcriptomic data of caste-specific phenotypes for nine species of social wasps, representing a range of levels of social complexity in the transition to superorganismality (Fig. 1). Using machine-learning algorithms we exploited these datasets to determine whether there is a conserved genetic toolkit for social behaviour across the major transition from non-superorganismal (simple) to superorganismal (complex) species within the same lineage (Aim 1). We then further interrogate these data to identify whether there are any key discernible differences in the molecular bases of social behaviour in the simpler versus the more complex societies (Aim 2). Accordingly, we provide the first evidence of a conserved genetic toolkit across the spectrum of the major transition to sociality in wasps; we also reveal novel insights into the molecular patterns and processes at a key transitional point of the major transition from simple sociality to complex superorganismality.

**Fig. 1.**
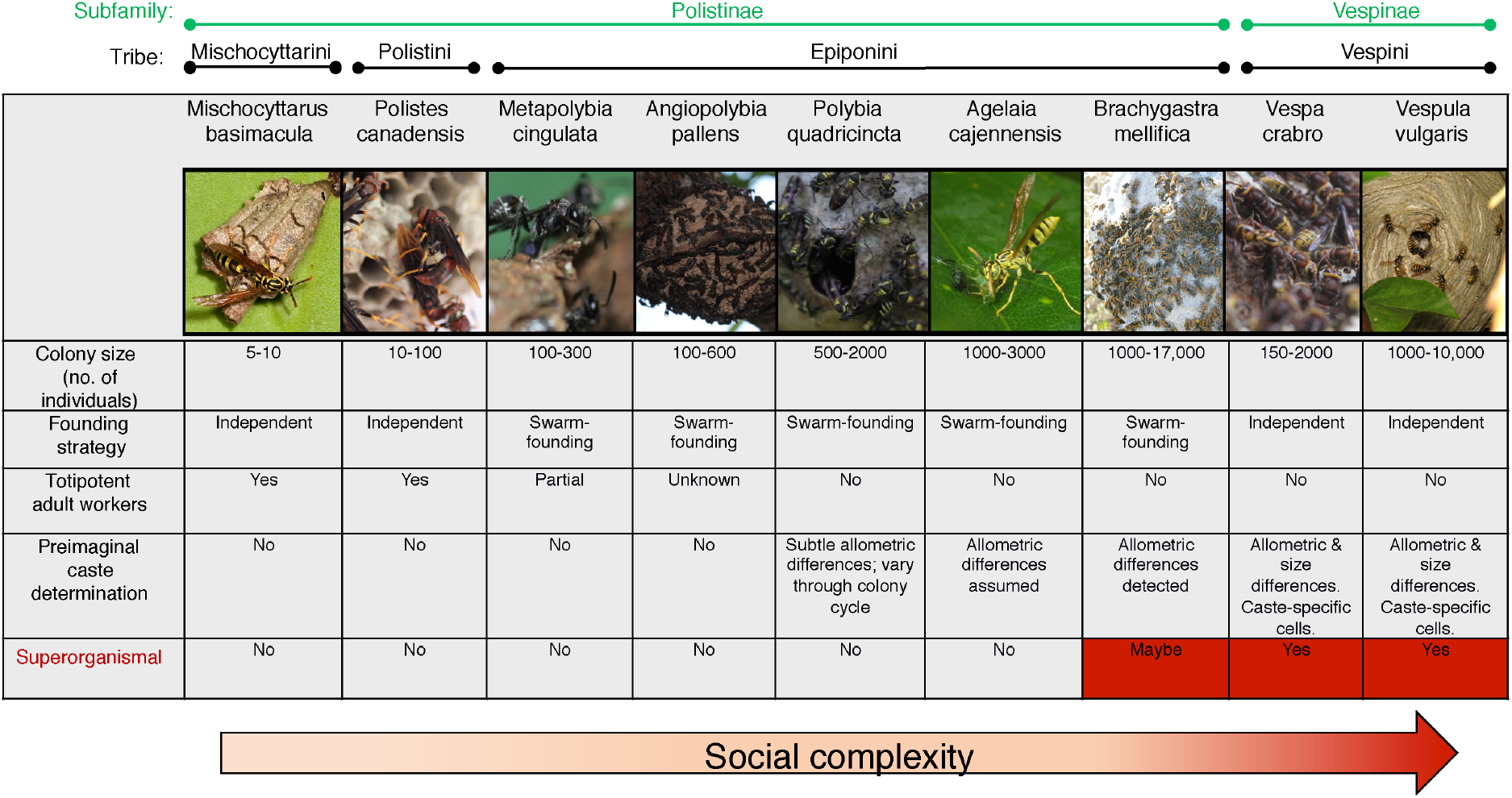
Social wasps as a model group. The nine species of social wasps used in this study, and their characteristics of social complexity. The Polistinae and Vespinae are two subfamilies comprising 1100+ and 67 species of social wasp respectively, all of which share the same common non-social ancestor, an eumenid-like solitary wasp^77^. The Polistinae are an especially useful subfamily for studying the process of the major transition as they include species that exhibit simple group living comprised of small groups (<10 individuals) of totipotent relatives, as well as species with varying degrees of more complex forms of sociality, with different colony sizes, levels of caste commitment and reproductive totipotency^78^. The Vespinae include the yellow-jackets and hornets, and are all superorganismal, meaning caste is determined during development in caste-specific brood cells; they also show species-level variation in complexity, in terms of colony size and other superorganismal traits (e.g. multiple mating, worker policing)^79^. Ranked in order of increasing levels of social complexity, from simple to more complex, these species are: *Mischocyttarus basimacula basimacula, Polistes canadensis, Metapolybia cingulata, Angiopolybia pallens, Polybia quadricincta, Agelaia cajennensis, Brachygastra mellifica, Vespa crabro* and *Vespula vulgaris* (see Supplementary Methods for further details of species choice). Where data on evidence of morphological castes was not available from the literature, we conducted morphometric analyses of representative queens and workers from several colonies per species. (see Supplementary Methods; Supplementary Table S1). Image credits: *M. basimacula* (Stephen Cresswell). *A. cajennesis* (Gionorossi; Creative Commons); *V. vulgaris* (Donald Hobern; Creative Commons). *V. crabro* (Patrick Kennedy); *P. canadensis; M. cingulata, A. pallens, P. quadricinta*, (Seirian Sumner), *B. mellifica* (Amante Darmanin; Creative Commons).

## Results

We chose one species from each of nine different genera of social wasps representing the full spectrum of social diversity within the Polistinae and Vespinae (see Figure 1; Supplemental Table S1). For each species, we sequenced RNA extracted from whole brains of adults to construct *de novo* brain transcriptomes for the two main social phenotypes – adult reproductives (defined as mated females with developed ovaries, henceforth referred to as ‘queens’ for simplicity; see Supplementary Table S2) and adult non-reproductives (defined as unmated females with no ovarian development, henceforth referred to as ‘workers’; see Supplementary Methods). Using these data we could reconstruct a phylogenetic tree of the Hymenoptera using single orthologous genes (Orthofinder^32^), resulting in expected patterns of phylogenetic relationships (Supp. Fig. 1). This dataset provides coverage across the spectrum of the major transition to sociality (see Fig. 1; Supplementary Table S1), and provides us with the opportunity to test the extent to which the same molecular processes underpin the evolution of social phenotypes across the spectrum of the major transition to superorganismality in wasps.

### Aim 1: Is there a shared genetic toolkit for caste among species across the major transition from non-superorganismality to superorganismality?

We found several lines of evidence of a shared genetic toolkit for caste across the wasp species using two different analytical approaches.

#### Caste explains gene expression variation, after species-normalisation

The main factor explaining individual-level gene expression variation was species identity (Fig. 2a). However, since we are interested in determining whether there is a shared toolkit of caste-biased gene expression across species, we needed to control for the effect of species in our data. To do this, we performed a between-species normalisation on the transcript per million (TPM) score, scaling the variation of gene expression to a range of −1 to 1 (see Supplementary Methods). After species-normalisation, the samples separate mostly into queen and worker phenotypes in the top two principle components (Fig. 2b). This suggests that subsets of genes (a potential toolkit) are shared across these species and are representative of caste differences. However, there were outliers: *Brachygastra* did not cluster with any of the other samples; *Agelaia* showed little caste-specific separation; and *Vespa* phenotypes clustered in the opposite direction to phenotypes in the other species. These initial data visualisations suggest that these species may not share the same caste-specific patterns as the other species, but we cannot rule out data and/or sampling anomalies, especially since gynes (unmated, newly-emerged queens) were included in the queen sample for *Vespa*.

**Fig. 2.**
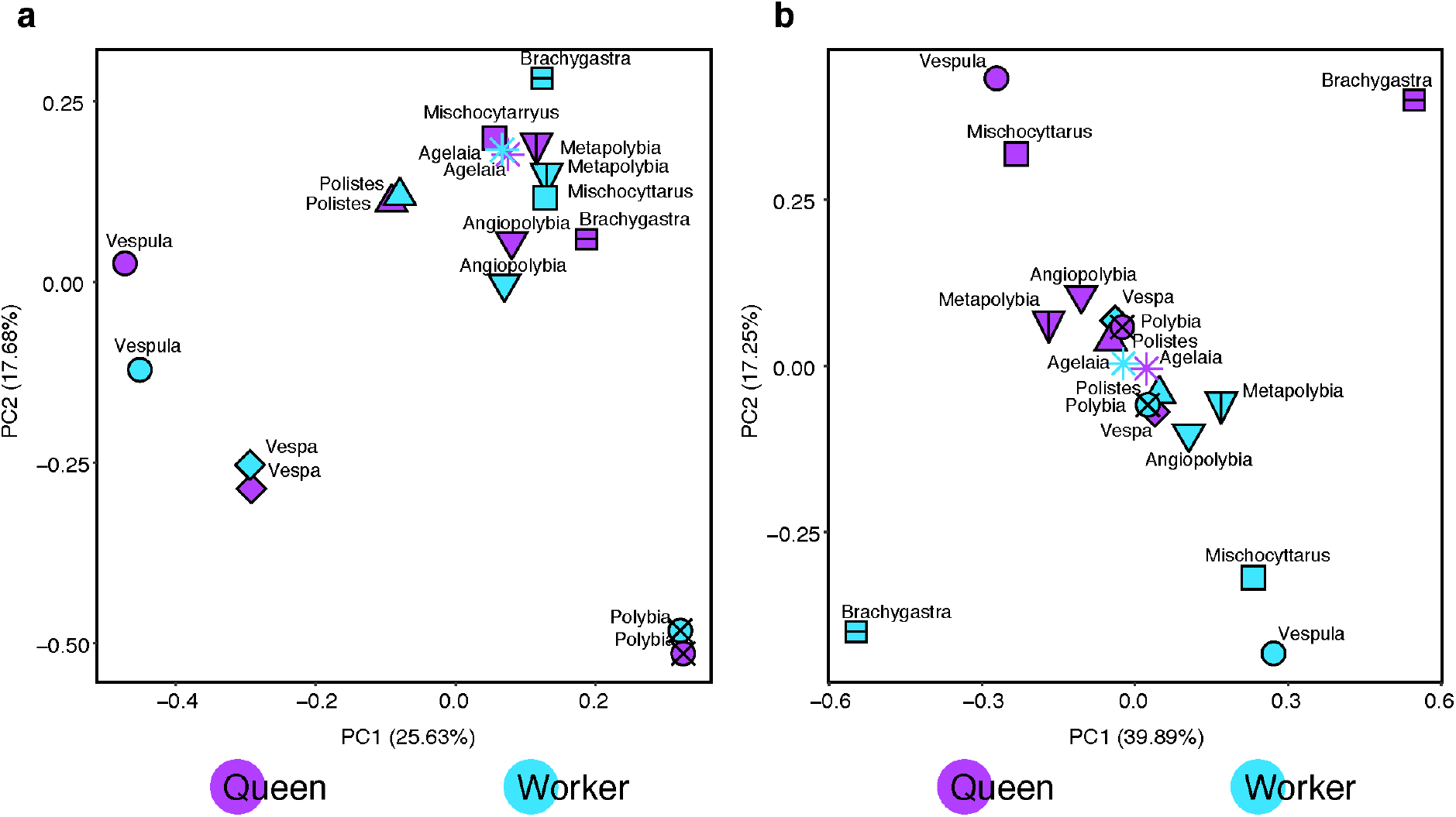
Principal component analyses of orthologous gene expression before and after between-species normalisation. **a)** Principal component analyses performed using log2 transcript per million (TPM) gene expression values. This analysis used single-copy orthologs (using Orthofinder), allowing up to three gene isoforms in a single species to be present, whereby we took the most highly expressed to represent the orthogroup, as well as filtering of orthogroups which have expression below 10 counts per million. **b)** Principal component analysis of the species-normalised and scaled TPM gene expression values using same filters as (a). Caste denoted by purple (queen) or blue (worker). Species are denoted by symbols.

Analyses of orthologous genes found in all nine species (Supplementary Table S3) revealed sets of caste-biased orthologous genes among the nine species; however, no orthologous DEGs showed consistent caste-biased differential expression across all nine wasp species (Fig. 3a; Supplementary Table S4; using unadjusted <0.05 p values). Depsite this, notable signatures of caste regulation were apparent, across the species set: e.g. orthogroup OG0002698 was differentially express across six of the nine species and is predicted to belong to the vitellogenin gene family (79.0% identity; using the *Metapolybia* protein sequence to represent the orthogroup), a well-known regulator of social behaviour in insects^33^. When the analysis was limited to caste-specific DEGs found in at least two species (n=95; Supplementary Table S4), there was overrepresentation of catabolic and metabolic GO terms (Fig. 3b; Supplementary Table S4).

**Fig. 3.**
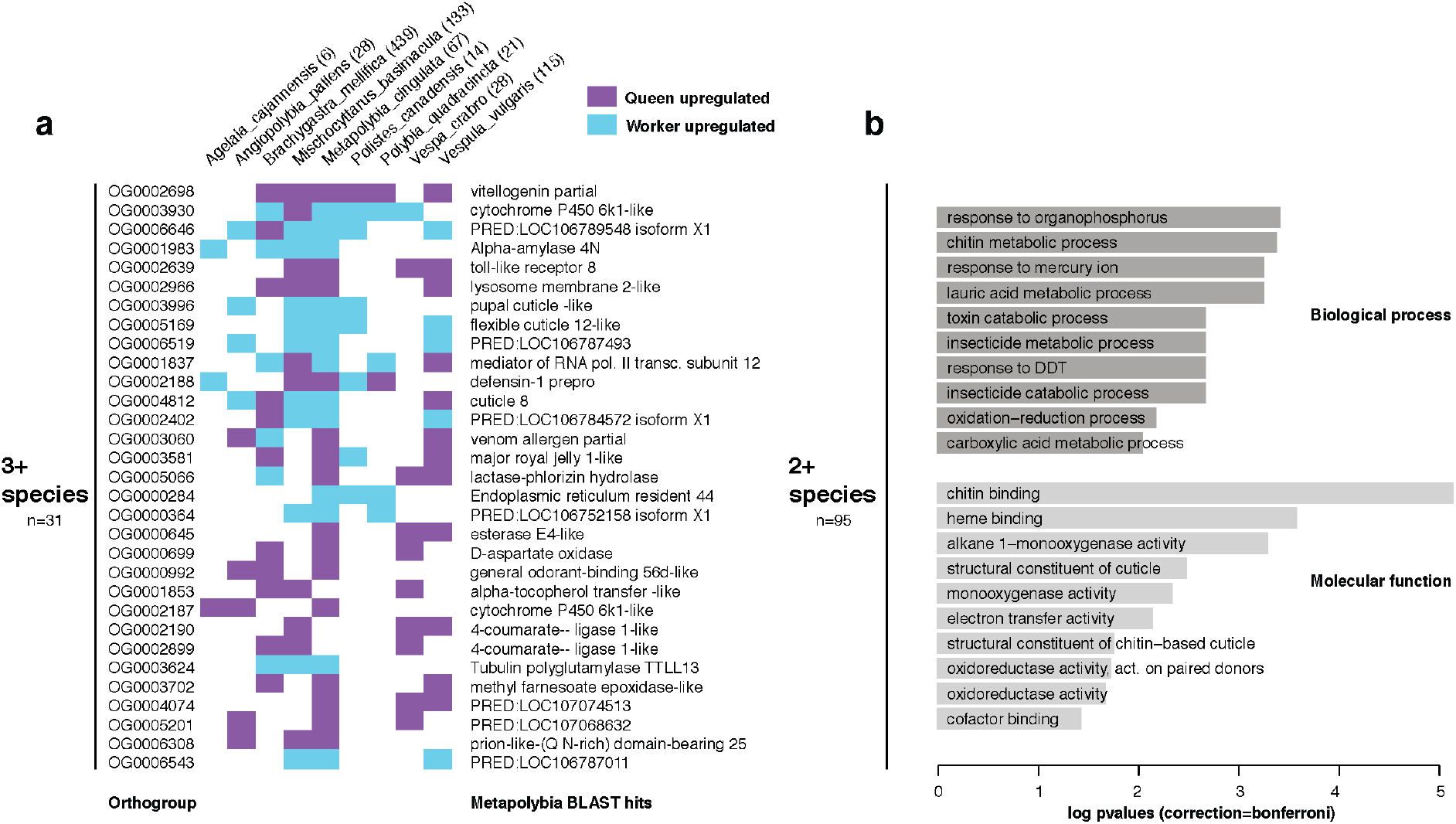
Overlap of differential caste-biased genes (queen vs worker) and their functions across eusocial wasp species. **a)** Heatmap showing the differential genes that are caste-biased in at least three species (identified using edgeR) using the orthologous genes present in the nine species. Listed for each species, is the total number of differentially expressed genes per species (orthologous-one copy only). *Metapolybia* Blast hits are listed. **b)** Gene ontology histogram of overrepresented terms of genes found differentially expressed in at least two out of the nine species (n=95 genes; in either queen or worker [not both]), with a background of those expressed in each species above 1 TPM. *P* values are single-tailed and were not corrected, given the low levels of enrichment generally and are therefore not significant for multiple testing.

#### A toolkit of many genes with small effects predicts caste across the spectrum of sociality in wasps

Conventional differential expression analyses (e.g. edgeR) require a balance of *P* value cut-offs and fold change requirements to reduce false-positive and false-negative errors^34^. Therefore, consistent patterns of many genes with smaller effect sizes may be missed when applying strict statistical measures. Support vector machine (SVM) learning approaches use a supervised learning model capable of detecting subtle but pervasive signals in differential expression between conditions (e.g. for classification of single cells^35,36^, cancer cells^37^ and in social insect castes^38^). We used this approach to test whether gene expression can successfully classify caste identity for unknown samples; accurate classification of samples as queens or workers based on their global transcription patterns would be evidence for a genetic toolkit underpinning social phenotypes.

Starting with a “leave-one-out” SVM approach, we attempted to classify samples of a test species as queens or workers, using a predictive gene set generated from a model trained on caste-specific gene expression from eight of the nine species, with the ninth species being the test sample. The analysis was repeated until each of the species had been ‘left out’ and their caste classification tested. Using 3486 single copy orthologues, and removing orthogroups with low expression (n=2020), we could filter the matrix by progressive feature selection (based on linear regression, to refine the gene sets to those that are informative; see Suppl. Methods), which reduces the number of genes used in the SVM, focussing on those genes informative for caste. When testing each left-out species, we largely attain accurate caste predictions for seven of the nine species, across most feature selection filters (Fig. 4a; > 0.5 likelihood in queen sample); the same two outlier species from Fig. 2 (*Agelaia* and *Brachygastra*) showed generally lower predictions of queen likelihood for the queen sample (<0.5). This suggests that many hundreds of genes may be caste-biased to some degree.

**Fig. 4.**
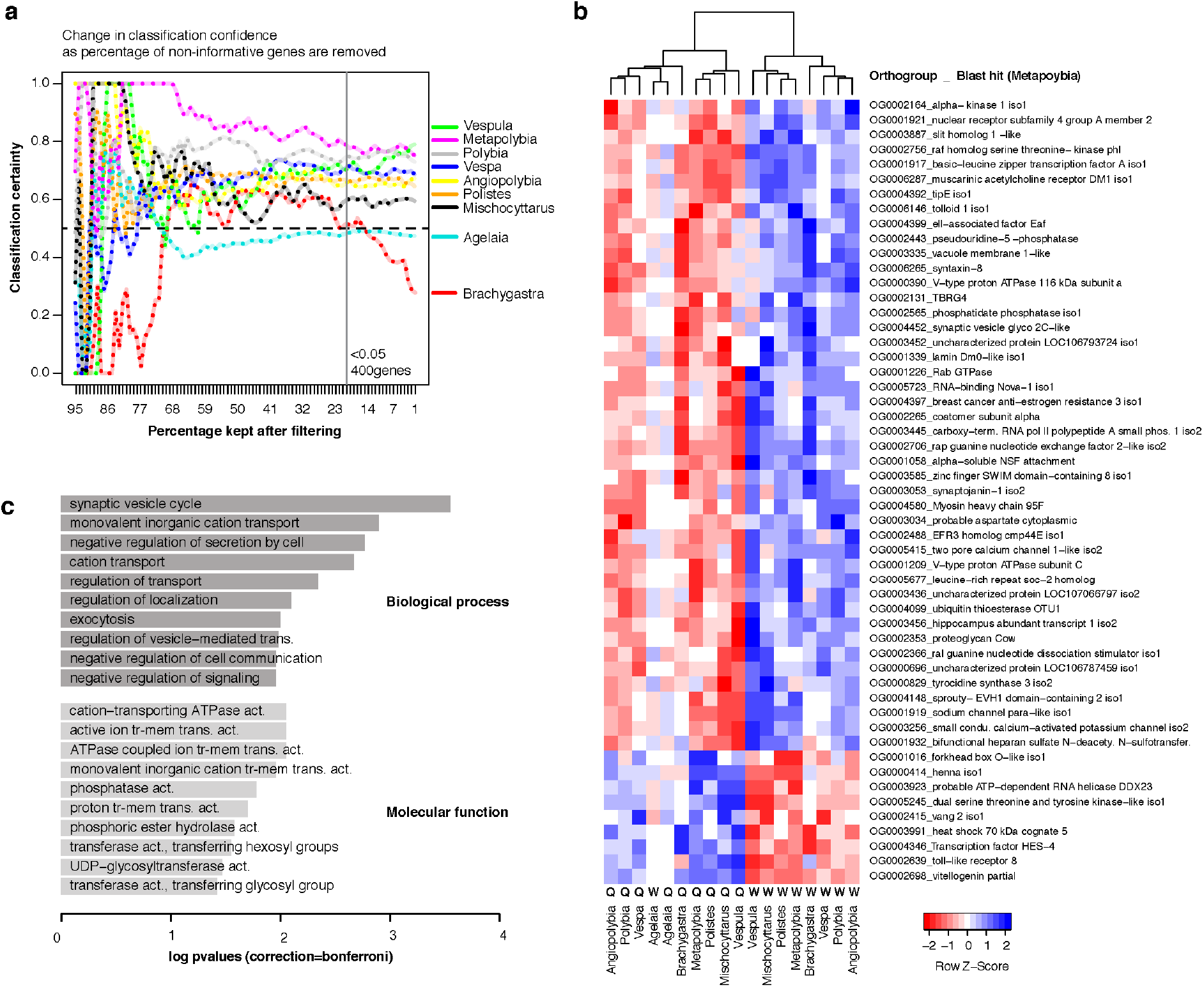
A genetic toolkit for social behaviours across eusocial wasps. **a)** Change in certainty of correct classifications through progressive feature selection. Models were trained on eight species and tested on the ninth species. Features (a.k.a. genes) were sorted by linear regression with regard to caste identity, beginning at 95% where almost all genes were used for the predictions of caste, to 1% where only the top one percent of genes from the linear regression (sorted by *P* value) were used to train the model. ‘1’ equates to high classification certainty. **b)** Heatmap of the top 53 species-normalised gene expression levels in the nine species with queen/worker indicated. Genes selected using linear regression (*P* value < 0.001) used in the SVM model, showing orthogroup name and top *Metapolybia* BLAST hit. **c**) Gene Ontology for the top 400 orthologous genes predictive of caste across species (linear regression *P* value < 0.05), and a background of all genes used in the SVM model (i.e. with a single gene representative for all species in the test). *P* values are bonferroni corrected and single-tailed. Abbreviations, “tr-mem”:transmembrane, “act.”: activity, “trans.”:transporter.

Within the SVM model of nine species (Fig. 4), we found 400 significant orthologs (genes) after feature selection with a *P* value of less than 0.05 (Supplementary Table S5; top 53 genes (p <0.001) shown in Fig. 4b). These 400 genes were enriched for neural vesicle transport related signalling functions (Fig. 4c; Supplementary Table S5), and may form the most important constituents of a shared toolkit for social behaviour across non-superorganismal and super-organismal social wasps.

Using Gene Set Enrichment Analysis (GSEA), we could compare the genes discovered in the two methods (edgeR and SVM), finding enrichment in the gene sets identified as important for social behaviour (Supplementary Figure 2). However, only ten genes were identified as significantly caste-biased in both methods (Supplementary Table S6). Of these, some have previously been identified as having relevance to social evolution and caste differentiation; these include Vitellogenin (mentioned earlier; OG0002698) and Cytochrome P450 (OG0000434)^10,39^, thought to be involved in chemical signalling between castes and associated with expression of juvenile hormone^39^. Further, UDP-glucuronosyltransferase 2C-like (OG0001554), downregulated in virgin versus mated fire ant queens^40^; esterase E4-like (OG0000645) upregulated in young honeybee queens compared to nurses at the proteomic level^41^; neprilsin-1 (OG0004128) is differentially expressed in major/minor *C. floridanus* workers and after caste reprogramming^42^, which could be involved in caste memory formation^43^. There are also other genes of interest, which to our knowledge have not previously associated with caste, including Toll-like receptor 8 (OG0002639) (see Supplementary Table S6).

### Aim 2: Are there different fine-scale toolkits that reflect different levels of social complexity?

To explore differential patterns *within* the conserved predictive toolkit for caste differentiation identified in Aim 1, we trained an SVM model using the four species with the simplest societies (*Mischocyttarus, Polistes, Metapolybia* and *Angiopolybia*) as representatives of the earlier stages in the major transition (see Fig. 1), and tested how well this gene set classified castes for the four species with the more complex societies as representatives of the later and superorganismal stages of the major transition (*Polybia, Agelaia, Vespa* and *Vespula;* see Fig. 1). *Brachygastra* was excluded due to its poor performance overall (see Fig. 4) and to ensure we compared the same number of training sets in each case. If castes in the test species classify well, this would suggest that the processes regulating castes in the simpler societies are also important in the more complex societies (i.e. there is no specific toolkit for simple sociality, which is then lost in the evolution of social complexity). Conversely, if the test species do not classify well, this would suggest that there are distinct processes regulating caste in the simpler societies that are lost (or become less important) in the evolution of more complex forms of sociality.

The putative toolkit for castes in the simplest societies consisted of ~1021 genes after feature selection (Supplementary Table S7 [Simple]). *Vespula* and *Polybia queens* classified extremely well (Fig. 5-upper); importantly, classifications for both these species improved with progressive feature selection. *Vespa* classified correctly but less well (likely because the queens included gynes); *Agelaia* classified to the wrong caste (consistent with results from Aim 1). Overall, based on these species, these results suggest that the genetic toolkit for simple societies is well conserved in the more complex societies that we sampled.

**Fig. 5.**
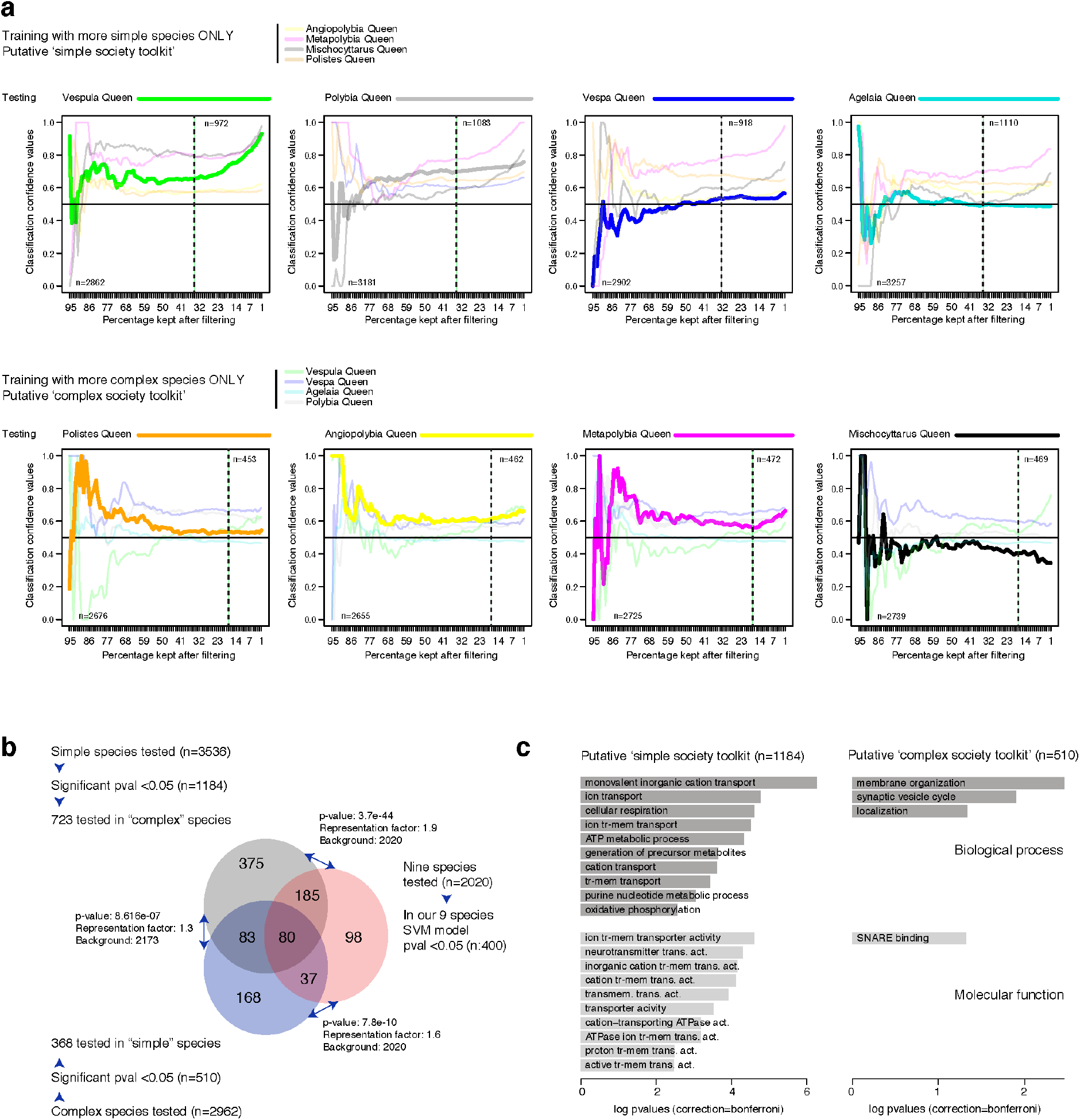
Testing for the presence of a defined ‘simple society toolkit’ and a ‘complex society toolkit’. **a)**: Using the four species with the more complex societies, or the four with the more simple societies, we trained and tested an SVM model, using progressive filtering of genes (based on the linear regression). Likelihood of being a queen from zero to 1 is plotted for each species across the progressively filtered sets. The number of genes used in the SVM model are shown for each test (bottom left of each panel), of which the total number of genes left after the regression filter are shown (top right of each panel), using genes with a *P* value < 0.05. For each test (pair of Queen/Worker in each species), the SVM model was run using genes with only 1 homologous gene copy per species (maximum of 3 isoforms merged). **b)** Overlap of significant genes in the different sets, compared to the 400 found using all nine species. For each experiment, the number of genes (orthogroups) tested is listed, then the number of genes significant after linear regression, and finally the number of genes that were also tested in the other two experiments. Significant overlap is shown using hypergeometric tests (one-tailed). Blue represents genes expressed in the four species with the most complex societies; grey those expressed in the four species with the most simple societies; pink are those expressed across all nine species. **c)** Enriched gene ontology terms (TopGO) using a background of all genes tested in each individual experiment, using a bonferroni corrected single-tailed *P* values.

We next conducted the reciprocal analysis, training the SVM using the four species with the more complex societies (*Polybia, Agelaia, Vespa* and *Vespula*) and testing it on the four species with simpler societies (*Mischocyttarus, Polistes, Metapolybia* and *Angiopolybia*). The toolkit for castes in these more complex societies was much smaller than the one for simple societies, consisting of ~464 genes after feature selection (Supplementary Table 7 [Complex]), possibly due to the greater taxonomic distances involved (inc. Polistinae and Vespinae). This putative toolkit for castes in more complex societies was less successful in classifying castes for the simpler societies (Fig. 5a-lower), than the reciprocal analysis (above; Fig. 5a-upper): although two species classified in the right direction (*Metapolybia* and *Angiopolybia*), their classifications have much lower confidence than in the reciprocal test; furthermore, for the two simplest societies, *Polistes* queens were classified close to 0.5 (meaning the gene sets were uncertain between queen/worker) and *Mischocyttarus* classified in the wrong direction (Fig. 5a-lower). These results raise the interesting idea that the processes regulating caste differentiation in species with more complex societies may be unimportant (or absent) in the simpler societies. In further support of this, the putative ‘simple society toolkit’ overlapped to a greater extent with the overall toolkit found across all species (Fig. 4) than those of the putative ‘complex society toolkit’ (Fig. 5b), hypergeometric overlap shown for both comparisons). Gene ontology results are similar between the two sets, and are composed of synaptic and membrane related terms (Fig. 5c; Supplementary Table 8); however the ‘simple society toolkit’ contains enrichment for metabolic/cellular respiration and ion/cation transport which are missing in the ‘complex society toolkit’.

We conducted additional tests to determine whether other factors could better explain the molecular basis of caste, besides level of social complexity, and to verify that our reciprocal SVM approach was valid given the small sample sizes. Using the same reciprocal SVM approach, we found that the molecular basis of a key life-history trait - nest founding behaviour - are largely conserved across species (Supplementary Figure 3; Supplementary Table 7[Swarm/Independent], Supplementary Table 8). From a biological perspective this suggests there is no specific genomic innovation associated with this life-history innovation that interacts with caste, as caste was correctly predicted in all species, with the exception of *Agelaia*. From a methodological perspective this indicates that the SVMs can perform well even using this small number of species unlikely to be affecting the performance of our social complexity SVM. Likewise, we tested for an effect of phylogeny, testing how well castes in the Vespines (*Vespa, Vespula*) classified using a putative Polistinae caste toolkit as the training set; there was little influence of subfamily on performance of the SVMs, with queens and workers being classified with 70-80% confidence (Supplementary Figure 3; Supplementary Table 7; the reverse of this test could not be performed due to low sample sizes for a Vespine training set). This suggests that the genes important for caste identity are shared across these two subfamilies.

## Discussion

Major transitions in evolution provide a conceptual framework for understanding the emergence of biological complexity. Discerning the processes by which such transitions arise provides us with critical insights into the origins and elaboration of the complexity of life. In this study we explored the evidence for two key hypotheses on the molecular bases of social evolution by analysing caste transcription in nine species of wasps. As predicted, we find evidence of a shared genetic toolkit across the spectrum of social complexity in wasps; importantly, using machine learning we reveal that this toolkit likely consists of many hundreds of genes of small effect (Fig. 4). However, in sub-setting the data by level of social complexity, two important new insights are revealed. Firstly, there appears to be a putative toolkit for castes in the simpler societies that largely persists across the major transition, through to superorganismality. Secondly, different (additional) processes appear to become important at more complex levels of sociality. Further sampling is required to determine the extent to which the role of these additional processes is driven by the evolution of superorganismality, and the point of no return in the major transition to sociality.

The first important finding is that we identified a substantial set of genes that consistently classify caste across most of the species, irrespective of the level of social complexity. The taxonomic range of samples used meant we were able to confirm that specific genes are consistently differentially expressed, with respect to caste, across the species. These patterns would be difficult to detect if only looking at a few species, species across several lineages, or species representing only a limited range of social complexity. In addition to typical caste-biased molecular processes, we also identified that genes related to synaptic vesicles are different between castes; this is interesting as the regulation of synaptic vesicles affects learning and memory in insects^44^. To our knowledge, this is the first evidence of what may be a conserved genetic toolkit for sociality, from the first stages of social living to true superorganismality, including intermediate stages of complexity, which putatively represent different points in the major transition. Greater taxonomic sampling will allow further exploration of how these genes and their regulation change across the major transition, and help recover the full spectrum of genes that may have been important in the evolution of sociality.

The underlying assumption, based on the conserved toolkit hypothesis, has been that whatever processes regulate castes in complex societies must also regulate castes in simpler societies. Unexpectedly, our analyses suggest there may be additional molecular processes underpinning castes that become important in the more complex levels of sociality. The predictive gene set identified in the SVM trained on more complex species performed less well in classifying caste than the predictive tool kit derived from the simpler species. There may be fundamental differences discriminating (near) superorganismal societies from non-superorganismal societies. This highlights the importance of examining different stages in the major transition when attempting to elucidate its patterns and processes.

There were two consistent outlier species in every stage of our analyses: *Agelaia* and *Brachygastra*. Although we cannot rule out issues with the data, all samples underwent the same rigorous QC testing at the lab, sequencing and bioinformatics stages and so are unlikely to fully explain these patterns. Another explanation is that they are genuinely biologically different to the other species. One of the most profound phenotypic innovations in social insect evolution is when caste becomes irreversibly committed during development^11,22,45^; this has been referred to as ‘the point of no return’ in evolutionary terms, as once a society is comprised of workers and queens who are mutually dependent on each other for colony function (like different cogs in the same machine), it is difficult to revert to independence^12^. After this point, the society can be considered as a definitive superorganism – with a new level of individuals and unit of selection^12^. Intriguingly, these two species are putatively at this point in the transition to super-organismality (Fig. 1). *Vespa* also failed to classify well in some analyses, but this is likely explained by the fact that the sample of queens included some gynes (unmated newly-emerged future-queens). Our morphometric analyses of *Brachygastra* (Supplementary Table S1) detected possible evidence of pre-imaginal caste determination, suggesting it is on the cusp of becoming superorganismal. Similarly, subtle differences in morphology among queens and workers of *Agelaia* suggests they too may have some level of pre-imaginal caste determination^46^. We speculate that the evolution of irreversible caste commitment (in superorganisms) is accompanied by a fundamental shift in the underlying regulatory molecular machinery such that species undergoing the transition to superorganismality may have to rewire the core set of genes involved in regulating caste.

Despite being able to extract consistent SVM predictions, our models are only as good as the initial data used to train them. Our study suffers from a few limitations. Firstly, the sample size (number of species) is relatively low; SVMs are generally used on very large datasets such as clinical trials in the medical sciences^47^. Although our models did perform well, the analyses would be more robust by using more species in the training datasets. Indeed, we observed reduced performance in our model predictions when fewer species were included in the training set. Secondly, by comparing across multiple species, we can only train our model on genes that have a single representative isoform per species in each separate test. This reduces the numbers of genes we can test in each SVM model, especially where more distantly related species are included. We overcame this limitation by merging gene isoforms within the same orthogroup (potential gene duplications), yet this comes with some additional costs as some genes are discarded in this process. Finally, genomes are not available for most of the species we tested; our measurements are based on *de novo* sequenced transcriptomes, which potentially contain misassembled transcripts, which could reduce the ability to find single copy orthologs across species. For these reasons, the numbers of genes detected in our putative toolkit for sociality is likely to be conservative and modest (potentially by several fold). These limitations are likely to apply to many similar studies, due to the difficulty and expense of obtaining high quality genomic data for specific phenotypes for non-model organisms. Our study illustrates the power of SVMs in detecting large suites of genes with small effects, which largely differ from those identified from conventional differential expression analysis^38^. We advocate the use of the two methods in parallel: our conventional analyses suggested that metabolic genes appear to be responsible for the differences between castes, whereas the SVM genes were mostly enriched in neural vesicle transportation genes, which have not previously been connected with caste evolution. SVMs may therefore reveal new target for genes involved in the evolution of sociality. We anticipate that bioinformatic and machine learning approaches, as demonstrated here, may become a useful tool in a wide range of ecological and evolutionary studies on the molecular basis of phenotypic diversity.

In conclusion, our analysis of brain transcriptomes for castes of social wasps suggest that the molecular processes underpinning sociality are conserved throughout the major transition to superorganismality. However, additional processes may come into play in more complex societies, putatively driven by selection happening at the point-of-no-return, where societies transition to become committed superorganisms. Importantly, this suggests there may be fundamental differences in the molecular machinery that discriminates superorganismal societies from non-superorganismal societies. The evolution of irreversible caste commitment (in superorganisms) may require a fundamental shift in the underlying regulatory molecular machinery. Such shifts may be apparent in the evolution of sociality at other levels of biological organisation, such as the evolution of multicellularity, taking us a step closer to determining whether there is a unified process underpinning the major transitions in evolution.

## Acknowledgements

We would like to thank Laura Butters for her help with the morphometric analyses of *Brachygastra*, James Carpenter at the American Natural History Museum and Christopher K. Starr for confirming species identification, and the Bristol Genomics Facility for their assistance with library preparations and sequencing. This work was conducted under collection and export permits for Trinidad (Forestry Division Ministry of Agriculture: #001162) and Panama (Autoridad nacional del ambiente (ANAM) SE/A-55-13). It was funded by the Natural Environment Research Council (NE/M012913/2; NE/K011316/1) awarded to SS, and a NERC studentship and Smithsonian Tropical Research Institute pre-doctoral fellowship awarded to E.F.B.

## Methods

### Study Species

Nine species of vespid wasps were chosen to represent different levels of social complexity across the major transition (Fig. 1). The simplest societies in our study are represented by *Mischocyttarus basimacula basimacula* (Cameron) and *Polistes canadensis;* wasps in these two genera are all independent nest founders and lack morphological castes (defined as allometric differences in body shape, rather than overall size) or any documented form of life-time caste-role commitment^48–51^. They live in small family groups of reproductively totipotent females, one of whom usually dominates reproduction (the queen); if the queen dies she is succeeded by a previously-working individual^21^. As such, these societies represent some of the earliest stages in the major transition, where caste roles are least well defined, and where individual-level plasticity is advantageous for maximising inclusive fitness.

The Neotropical swarm-founding wasps (Hymenoptera: Vespidae; Epiponini) include over 20 genera with at least 229 species, exhibiting a range of social complexity measures, from complete absence of morphological caste (pre-imaginal) determination to colony-stage specific morphological differentiation, through to permanent morphological queen-worker differentiation^52^. As examples of species for which there is little or no evidence of developmental (morphological) caste determination, we chose *Angiopolybia pallens* which is phylogenetically basal in the Epiponines^53,54^ and *Metapolybia cingulata* (Fabricius)^53,54^. We confirmed the lack of clear caste allometric differences in *M. cingulata* as data were lacking (see Morphometrics methods (below) and Supplementary File S1).

As examples of species showing subtle, colony-stage-specific caste allometry, we chose a species of *Polybia*. The social organisation of *Polybia* spp is highly variable, ranging from complete absence of morphological queen-worker differentiation^55^. *Polybia quadricincta* is a relatively rare (and little studied) epiponine wasp which can be found across Bolivia, Brazil, Columbia, French Guiana, Guyana, Peru, Suriname and Trinidad (Richards, 1978). Our morphometric analyses found some evidence of subtle allometric morphological differentiation in this species, but with variation through the colony cycle (Supplementary File S1); this suggest it is a representative species for the evolution of the first signs of pre-imaginal caste differentiation.

Many species of the genera *Agelaia* and *Brachygastra* appear to show pre-imaginal caste determination with allometric morphological differences between adult queens and workers^53^. We chose one species from each of these genera as representatives of the most socially complex Polistine wasps. Although no morphological data were available for *Agelaia cajennensis* (Fabricius) all species of *Agelaia* studied show some level of preimaginal caste determination^53,56^. *Brachygastra* exhibit a diversity of caste differentiation^53,57^; our morphological analysis of caste differentiation *B. mellifica* confirms that this species is highly socially complex, with large colony sizes^53^ and pre-imaginal caste determination resulting in allometric caste differences (Supplementary File S1).

All species of Vespines are independent nest founders and superorganisms, with a single mated queen establishing a new colony alone and with morphological castes that are determined during development. However, some species exhibit derived superorganismal traits, such as multiple mating^58^, which have likely evolved under different selection pressures to the major transition itself^59^. The European hornet, *Vespa crabro*, exhibits the hallmarks of superorganismality (see Fig. 1) but little evidence of more derived superorganismal traits, such as high levels of multiple mating. Conversely, multiple mating is common in *Vespula* species, including *V. vulgaris* with larger colony sizes than *Vespa^58^*, suggesting a more complex level of social organisation.

### Sample collection

Where possible, we sampled from colonies representing different stages in the colony cycle, as caste differentiation can vary as the colony matures in some species (Supplementary File S2). *Metapolybia cingulata* (6 colonies), *Polistes canadensis* (3 colonies), *Agelaia cajennensis* (1 colony) and *Mischocyttarus basimacula basimacula* (3 colonies) were collected from wild populations in Panama in June 2013. *Brachygastra mellifica* (4 colonies) were collected from populations in Texas, USA in June 2013. *Angiopolybia pallens* (2 colonies) and *Polybia quadricincta* (2 colonies) were collected from Arima Valley, Trinidad in July 2015. *Vespa crabro* (4 colonies) and *Vespula vulgaris* (4 colonies) were collected from various locations in South East England, UK in 2017. Queens and workers were collected directly from their nests during the daytime, placed immediately into *RNAlater* (Ambion, Invitrogen) and stored at −20°C until further use. An exception was that gynes (newly-emerged, unmated queens) in addition to queens were used for *V. crabro* due to difficulty in obtaining samples of mature queens. Samples were ultimately pooled *within castes* for bioinformatics analyses, such that each informatic pool consisted of 3-6 individual brains from wasps sampled across 2-4 colonies to capture individual-level and colony-level variation in gene expression (see Supplementary Table S2). Samples of *M. cingulata, A. cajennensis, M. basimacula and B. mellifica* were sent to James Carpenter at the American Natural History Museum for species verification. *A. pallens* and *P. quadricincta* were identified by Christopher K. Starr, at University of West Indies, Trinidad and Tobago.

### Morphometrics

Data on morphological differentiation among colony members (and thus information on whether pre-imaginal (developmental) caste determination was present) was lacking for *M. cingulata, P. quadricinta* and *B. mellifica;* therefore, we conducted morphometric analyses on these three species in order to ascertain the level of social complexity. Morphometric analyses were carried out using GXCAM-1.3 and GXCapture V8.0 (GT Vision) to provide images for assessing morphology. We measured 7 morphological characters using ImageJ v1.49 for queens and workers for each species. The body parts measured were: head length (HL), head width (HW), minimum interorbital distance (MID), mesoscutum length (MSL), mesoscutum width (MSW), mesosoma height (MSH) and alitrunk length (AL) (for measurement details, see ^60^). Abdominal measurements were not recorded as ovary development could alter the size of abdominal measurements, therefore biasing the results. The morphological data were analysed to determine whether the phenotypic classification, as determined from reproductive status, could be explained by morphological differences. ANOVA was used to determine size differences between castes for each morphological characteristic. A linear discriminant analysis was also employed to see if combinations of characters were helpful in discriminating between castes. The significance of Wilks’ lambda values were tested to determine which morphological characters were the most important for caste prediction. All statistical analyses were carried out using SPSS v23.0 or Exlstat 2018. Data and analyses given in Supplementary Table S1.

### Dissections & RNA extractions

Individual heads were stored in RNAlater for brain dissections; abdomens were removed and dissected to determine reproductive status. Ovary development was scored according to ^31,61^ and the presence/absence of sperm in the spermathecae was identified to determine insemination. Inseminated females with developed ovaries were scored as ‘queens’; non-inseminated females with undeveloped ovaries were scored as workers. Brains were dissected directly into RNAlater; RNA was extracted from individuals and then pooled after extraction into caste-specific pools; pooling after RNA extraction allowed for elimination of any samples with low quality RNA. Pooling individuals was generally necessary to ensure sufficient RNA for analyses, as well as accounting for individual variation to ensure expression differences are due to caste or species, and not dependent on colony or random differences between individuals. One exception to this was the *V. vulgaris* samples which were sequenced as individual brains and pooled bioinformatically after sequencing. Individual sample sizes per species are given in Supplementary Table S2.

Total RNA was extracted using the RNeasy Universal Plus Mini kit (Qiagen, #73404), according to the manufacturer’s instructions, with an extra freeze-thawing step after homogenization to ensure complete lysis of tissue, as well as an additional elution step to increase RNA concentration. RNA yield was determined using a NanoDrop ND-8000 (Thermo Fisher Scientific); all samples showed A260/A280 values between 1.9 and 2.1. An Agilent 2100 Bioanalyser was used to determine RNA integrity. Samples of sufficient quality and concentration were pooled and sent for sequencing. Libraries were prepared using Illumina TruSeq RNAseq sample prep kit at the University of Bristol Genomics Facility. Five samples were pooled per lane to give ~ 50M read per sample. Paired-end libraries were sequenced using an Illumina HiSeq 2000. Descriptions of pooling of individuals and pooled sets into single representatives of caste are shown in (Supplementary table S2). Raw reads are available on SRA/GEO (GSE159973).

### Preparation of *de novo* transcriptomes

Transcriptomes of *Agelaia, Angiopolybia, Metapolybia, Brachygastra, Polybia, Polistes* and *Mischocyttarus* were assembled using the following steps. First, reads were first filtered for rRNA contaminants using tools from the BBTools (version:BBMap_38) software suite (https://jgi.doe.gov/data-and-tools/bbtools/). We then used Trimmomatic v0.39^62^ to trim reads containing adapters and low-quality regions. Using these filtered RNA sequences, we could assemble a *de novo* transcriptome for each species (merging queen and worker samples) using Trinity v2.8^63^ and filter protein coding genes to retain a single transcript (most expressed) for each gene and transcript per million value (TPM), which we use for the rest of the analyses.

For *Vespula* and *Vespa*, reads from both queen and worker samples were assembled into *de novo* transcriptomes using a Nextflow pipeline (github:biocorecrg/transcriptome_assembly). This involved read adapter trimming with Skewer^64^, *de-novo* transcriptome assembly with Trinity v2.8.4^63^ and use of TransDecoder v5.5.0^63^ to identify likely protein-coding transcripts, and retain all translated transcripts. These were further filtered to retain the largest open read frame-containing transcript, which we listed as the major isoform of each protein. Trinity assembly statistics are shown in Supplementary Table S2.

### Measuring gene expression within-species

We calculated abundances of transcripts within queen and worker samples using “align_and_estimate_abundance.pl” within Trinity, using estimation method RSEM v1.3.1^65^, “trinity_mode” and bowtie2^66^ aligner. We then used edgeR v3.26.5 ^67^ (R version 3.6.0) to compare gene expression between queens and workers. Because we were comparing a single sequencing pool of several individuals per caste, we used a hard-coded dispersion of 0.1 and the robust parameter set to true to account for n = 1. Raw *P* values for each gene were corrected for multiple testing using a false discovery rate (FDR) cut-off value of 0.05.

We did not take advantage of genome data (where available), as only two of the species had published genomes at the time of analysis; using transcriptome-only analyses makes the analysis more consistent across species. Trinity assemblies and RSEM counts are available on GEO/SRA (GSE159973)

### Identification of orthologs

To identify gene-level orthologs, we used Orthofinder v.2.2.7^68^ with diamond blast^32,69^, multiple sequence alignment program MSA^65^ and tree inference using FastTree v2.1.10^70^. for our focal nine species, plus four out-group Hymenoptera species (Supplementary Fig. 1) and *Drosophila melanogaster*. The largest spliced isoform per gene (from Trinity) was designated the representative sequence for each gene. For subsequent analyses using the orthofinder table of genes, we allowed the merging of genes belonging to the same species in a single orthogroup (potential duplications). This decision has consequences for the number of genes we can use to test in our models, as the more species used will reduce the numbers of genes (with 1 to 1 orthology across all the species used in Orthofinder and SVM models). In order to get a sufficient number of single-copy gene orthogroups, we merged the genes in one species where there were three or less representative isoforms, only keeping the gene most highly expressed.

### Comparing gene expression between species

To compare gene expression between species, we focused on our set of shared one-to-one orthologs (merging 3 or less isoforms per species). We began by computing log transformed TPMs (transcripts per million reads) for each gene in each sample from the raw counts, followed by quantile normalisation. Next, we normalised for species, using an approach that is comparable to calculating a species-specific z-score for each sample. Specifically, we transformed the expression scores calculated above by subtracting the species mean and dividing by the species mean for each sample within a species. This calculation has two important effects. Firstly, subtracting the species mean from each sample within a species centres the mean expression of each species on zero, making the units of expression more comparable across species. Secondly, dividing by the species mean from each sample standardises the expression scores, producing a measure that is independent of the units of measurement, so that the magnitude of difference between queens and workers in each sample is no longer important. The transformed expression score thus allows us to focus on the relative expression in queens versus workers across species. Finally, we removed orthogroups where the counts per million were below 10 in both Queen and Worker samples of each species, to remove lowly expressed genes that may contribute noise to subsequent analyses. We then performed principal component analysis (PCA) in R on the raw TPM values and those with species scaling.

### Machine learning (support vector machines)

Support vector machines (SVM) were used to classify caste across the species. In brief, starting with a matrix of gene expression values, we performed pre-filtering steps (feature selection), before training a model and testing this on an additional dataset. The code to run these steps is shown on github (https://github.com/Sumner-lab/Multispecies_paper_ML). In summary, this involved taking species-scaled, logged and normalised matrix (from RSEM), with filtering of lowly expressed genes (as above), then invoking SVM predictions (radial model) and plotting; code was executed in perl or R. The full detail of these steps are outlined below.

To perform feature selection we identified only those orthologs that showed some association with caste across species. For this we used linear regression of each gene on caste: lm(caste ~ expr, data), using the training data only. With regression beta coefficients per orthologous gene, we could then rank genes by their statistical association with caste (Supp. Table 4 using the absolute values of the regression coefficients). This enabled us to measure how the classification certainty changed as we filtered out genes statistically un-associated with reproductive division of labour (Figure 4a). This basic feature selection approach is widely used to filter large datasets in the machine learning models ^71^.

Classification certainty of 0.5 would indicate the SVM could not tell the difference between the two castes (maximal uncertainty), and a classification certainty of 0/1 (worker or queen) would indicate that the SVM could predict caste accurately every time (maximal certainty).

After identifying candidate toolkit genes of reproductive division of labour, we tested whether or not they could be used to predict caste in unseen data. To do this, we trained support vector machines (SVM) using the R package e1071^72^. Radial kernels were chosen for the svm, which had better error statistics. We used a “leave-one-out” cross validation procedure to see how well an SVM could predict the castes of our samples, where the model is trained on all but one species and tested on the removed species.

### GO/GSEA enrichment and BLAST

To perform GO enrichment tests, we used the R package TopGO v2.42.0^73^, using Bonferroni cut-off *P* values of <=0.05. In order to assign gene ontology terms to genes in our new species, we used our Orthofinder homology table with annotations to *Drosophila melanogaster* (downloaded from Ensembl Biomart 1.10.2019). Within species, we calculated enrichment of each species’ gene to a background of all the genes expressed above a mean of 1 TPM. When comparing across the orthogroups (OG), we used *Metapolybia* GO annotations (derived from homology to *Drosophila*), with a background of those genes that have a mean >1 TPM in all species orthologues. GO comparisons were similar using other species as a database of gene to GO terms.

Using default settings in GSEA v4.0.3^74^ we compared the lists derived from the SVM experiment and conventional differential expression analysis (using the preranked mode). First (Supplementary Figure 2a), using the list of 2020 SVM (9 species) orthologs (excluding low-expression genes) ranked from 1 to 2020 based on the linear regression *P* values we could derive enrichment scores from the DEGs (n=95; from edgeR), where the total were reduced to 19 genes that were present in both analyses. Second (Supplementary Figure 2b), we ranked Vespula (Trinity) differentially expressed genes by log fold change, deriving enrichment scores with the 400 SVM genes significant in the nine species SVM, after linear regression p.value cutoff of 0.05. Blast2GO v 1.4.4^75^ using Metapolybia gene sequences using was used to annotate sequences, along with manual use of NCBI blastn^76^ suite online.

## Abbreviations

DOL: division of labour
ORF: open reading frame
MT: major transition
GO: gene ontology
SRA: sequence read archive
NCBI: National Centre for Biotechnology Information
BUSCO: Benchmarking set of Universal Single-Copy Orthologs
SVM: support vector machine
PCA: principal components analysis
Blast: Basic local alignment search tool
nr: non-redundant

## Author’s contributions

SS conceived the study and supervised the project; SS, EL, EB and BT collected the samples; DT, EB, BT and RB performed molecular lab work; DT & RB carried out the morphological work; MB & CW executed the bioinformatics pipelines, performed the statistical analyses; CW & SS drafted the manuscript, with input from all authors.

## Supplementary Figures

**Supp. Figure. 1.**
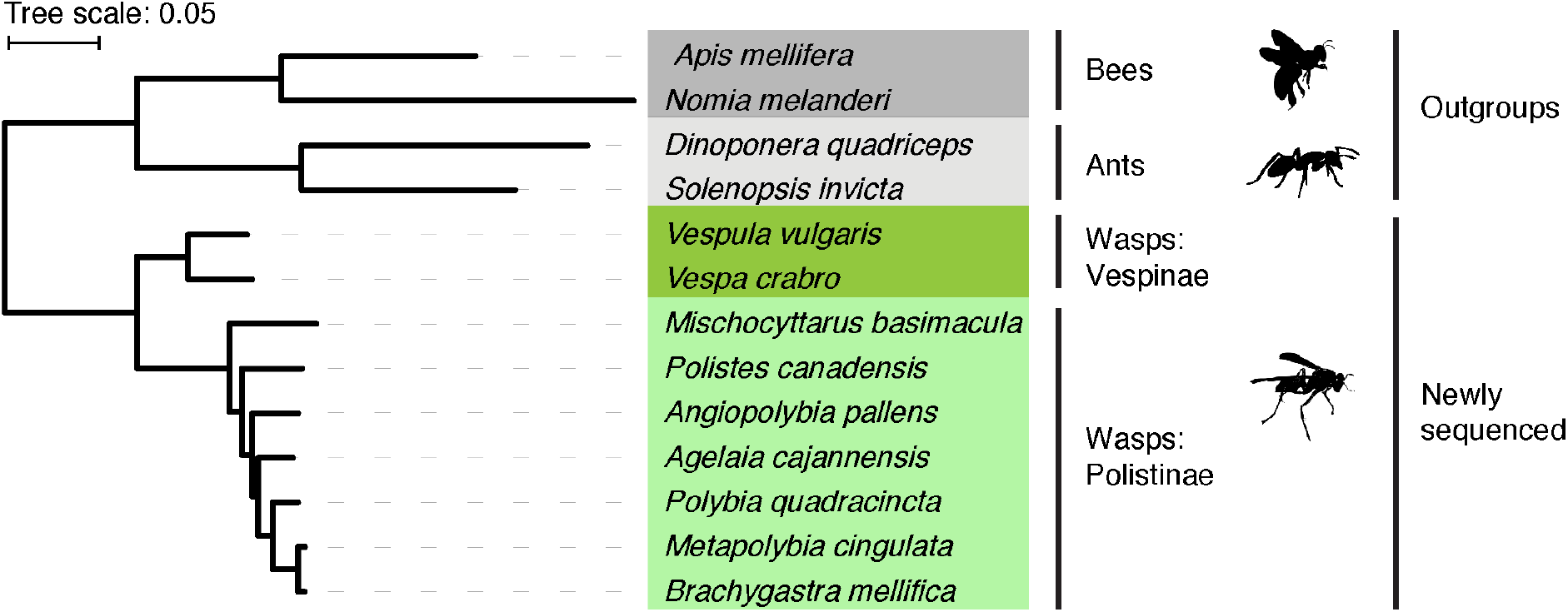
Phylogenetic tree of social wasps used in this study, and related hymenopterans generated using Orthofinder (SpeciesTree_rooted.txt). *Drosophila* was chosen as the root of the species tree (not shown), with two representative ants (*Dinoponera quadriceps* and *Solenopsis invicta*) and two bees (*Apis mellifera* and *Nomia melanderi*). Colours show groupings of ants, bees and wasps (Vespinae or Polistinae). For the nine wasp species (this study) we have sequenced adult caste-specific brain transcriptomic data (queen and worker). As expected: Vespinae are clearly separated from the Polistinae; *Angiopolybia* is basal to the other Epiponini; independent-founding, non-superorganismal wasps (*Polistes* and *Mischocyttarus*) are basal to the Polistinae^77,80^.

**Supp. Figure. 2.**
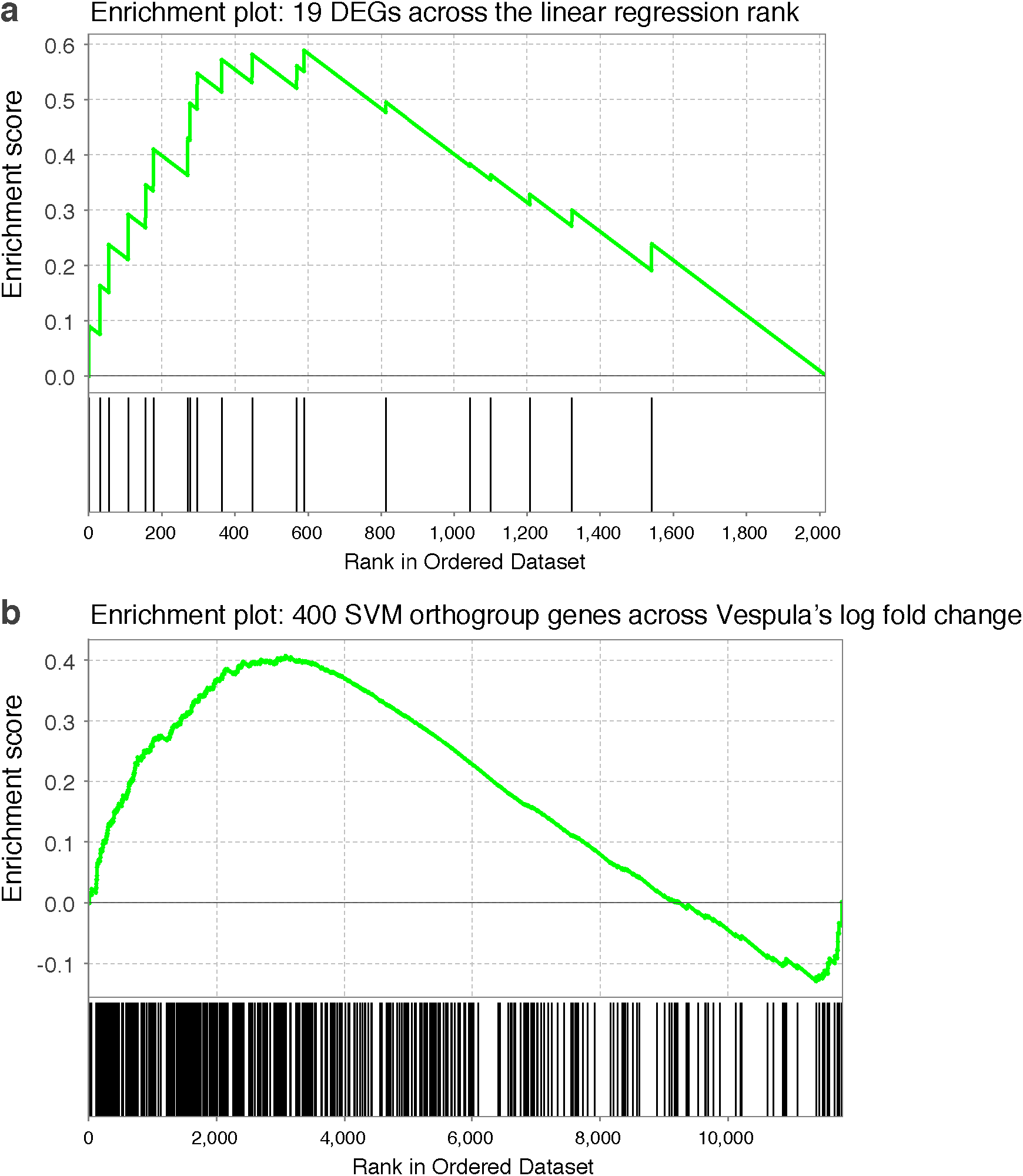
GSEA –Gene Set Enrichment Analysis comparing overlap of the orthogroups discovered with differentially expressed genes. **a):** Enrichment scores using the 95 orthogroups differentially expressed in at least two wasp species and their ordering across the ranked SVM genes derived from linear regression (2020 orthogroups; where rank 0 is the orthogroup most associated with caste). Of 95 genes, only 19 orthogroups were found in the linear regression sorted set. The upper plot shows the enrichment scores, while the lower plot shows the position of the 19 orthogroup genes across the ranked SVM list. Enrichment was found toward the higher ranks (nearer 0), suggesting that there is some overlap in genes found using linear regression (SVM)/edgeR approaches. **b):** Enrichment scores using the 400 significant SVM genes (from linear regression, converted to Vepsula IDs) across the log fold changes of Vespula genes from edgeR. Enrichment in both queen and worker biased ends (see two peaks: left/right) were detected, again suggesting limited overlap in the genes found using the two approaches.

**Supp. Figure. 3.**
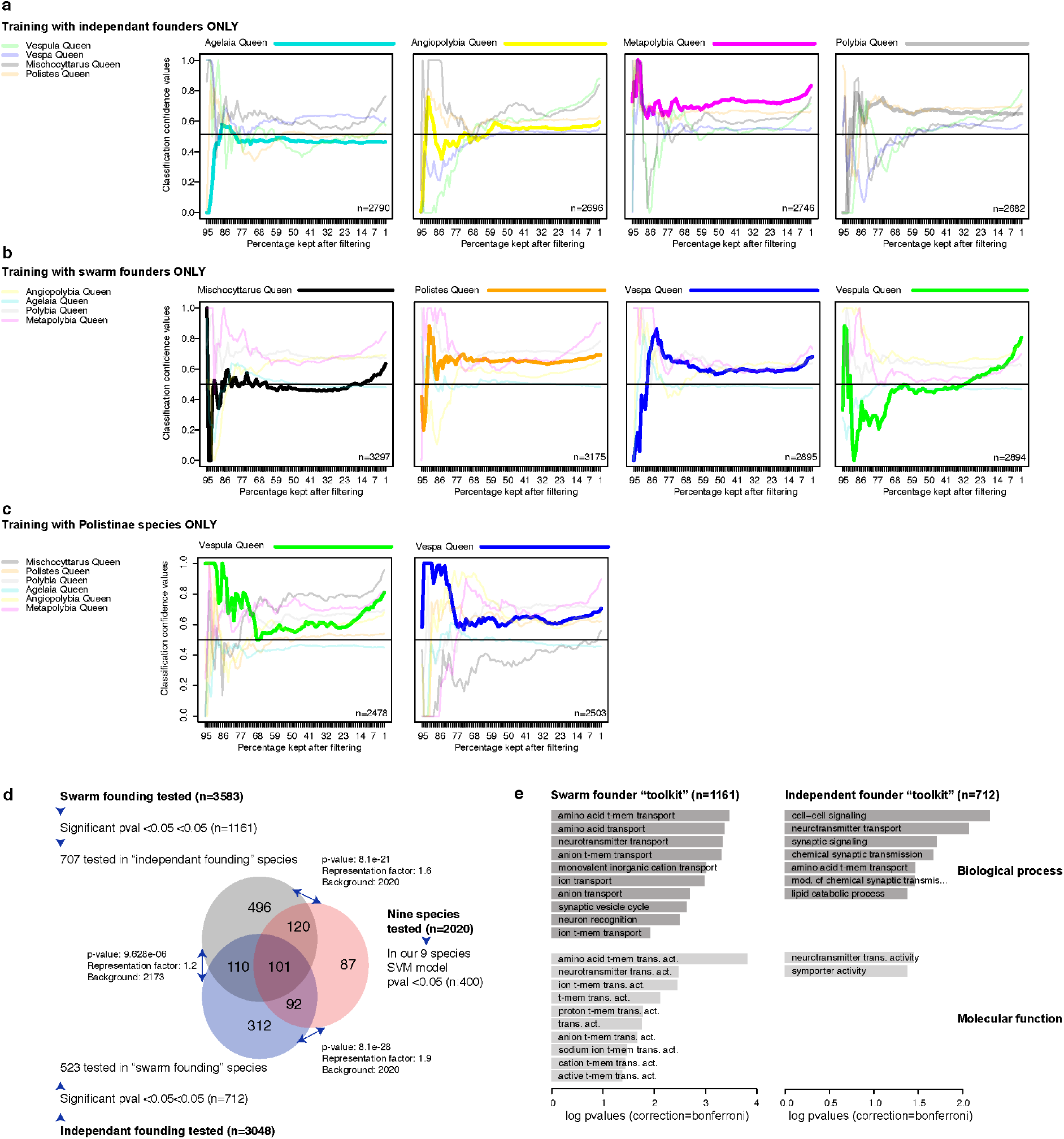
SVM predictions using founding behaviour and phylogeny to subset the training data. SVM predictions are given for each single species given training on species listed to left, showing the prediction after progressive feature selection from 95 to 1% of genes remaining after selection. Numbers of genes in each test are indicated in the lower left part of each plot. **a)** Training with independent-founders only, tested on the four swarm-founders (see Figure 1 for species). **b)** The reverse of ‘a’. **c)** Training with Polistine species only, and testing on the Vespines (*Vespa* and *Vespula*). The reverse was not possible, as the SVM requires a minimum number of samples to work. **d)** Overlap of genes found using the swarm/independent founding toolkits, along with the overlap with the 400 genes discovered using all nine species. Hypergeometric p.values and representation factors are shown. For each set we show the total number of genes tested in each experiment, followed by the number significant at the <0.05 pvalue cutoff, and then of these, the number that were also tested in the other two comparisons. These genes numbers are then overlapped in the Venn. **e)** Enriched gene ontology terms (TopGO) using a background of all genes tested in each individual experiment, using a bonferroni corrected single-tailed *P* values.

## References

1. Szathmáry, E. & Maynard Smith, J. The major evolutionary transitions. Nature 374, 227–232 (1995).

2. Kennedy, P. et al. Deconstructing Superorganisms and Societies to Address Big Questions in Biology. Trends in Ecology and Evolution 32, 861–872 (2017).

3. Toth, A.L. & Robinson, G. E. Evo-devo and the evolution of social behavior. Trends Genet. 23, 334–341 (2007).

4. Berens, a. J., Hunt, J. H. & Toth, a. L. Comparative Transcriptomics of Convergent Evolution: Different Genes but Conserved Pathways Underlie Caste Phenotypes across Lineages of Eusocial Insects. Mol. Biol. Evol. 32, 690–703 (2014).

5. Qiu, B. et al. Towards reconstructing the ancestral brain gene-network regulating caste differentiation in ants. Nat. Ecol. Evol. 2, 1782 (2018).

6. Warner, M. R., Qiu, L., Holmes, M. J., Mikheyev, A. S. & Linksvayer, T. A. Convergent eusocial evolution is based on a shared reproductive groundplan plus lineage-specific plastic genes. Nat. Commun. 10, 1–11 (2019).

7. Patalano, S. et al. Molecular signatures of plastic phenotypes in two eusocial insect species with simple societies. Proc. Natl. Acad. Sci. U. S. A. 112, 13970–5 (2015).

8. Toth, Amy L. Rehan, S.. Climbing the social ladder: The molecular evolution of sociality. (2015).

9. West-Eberhard MJ. Wasp societies as microcosms for the study of development and evolution. in Natural history and evolution of paper wasps. (eds. Turillazzi, S. & West-Eberhard, M. J.) 290–317 (Oxford University Press, 1996).

10. Toth, A. L. & Rehan, S. M. Molecular Evolution of Insect Sociality: An Eco-Evo-Devo Perspective. Annual Review of Entomology (2017). doi:10.1146/annurev-ento-031616-035601

11. Boomsma, J. J. Lifetime monogamy and the evolution of eusociality. Philos. Trans. R. Soc. Lond. B. Biol. Sci. 364, 3191–207 (2009).

12. Boomsma, J. J. & Gawne, R. Superorganismality and caste differentiation as points of no return: how the major evolutionary transitions were lost in translation. Biol. Rev. (2017). doi:10.1111/brv.12330

13. Ferreira, P. G. et al. Transcriptome analyses of primitively eusocial wasps reveal novel insights into the evolution of sociality and the origin of alternative phenotypes. Genome Biol. 14, R20 (2013).

14. Sumner, S. The importance of genomic novelty in social evolution. Mol. Ecol. 23, (2014).

15. Feldmeyer, B., Elsner, D. & Foitzik, S. Gene expression patterns associated with caste and reproductive status in ants: Worker-specific genes are more derived than queen-specific ones. Mol. Ecol. 23, 151–161 (2014).

16. Johnson, B. R. & Tsutsui, N. D. Taxonomically restricted genes are associated with the evolution of sociality in the honey bee. BMC Genomics 12, 164 (2011).

17. Simola, D. F. et al. Social insect genomes exhibit dramatic evolution in gene composition and regulation while preserving regulatory features linked to sociality. Genome Res. 23, 1235–1247 (2013).

18. Harpur, B. a et al Population genomics of the honey bee reveals strong signatures of positive selection on worker traits. Proc. Natl. Acad. Sci. U. S. A. 111, 2614–9 (2014).

19. Rubin, B. E. R., Jones, B. M., Hunt, B. G. & Kocher, S. D. Rate variation in the evolution of non-coding DNA associated with social evolution in bees. Philos. Trans. R. Soc. B Biol. Sci. 374, (2019).

20. Kapheim, K. M. Genomic sources of phenotypic novelty in the evolution of eusociality in insects. Curr. Opin. Insect Sci. 1–9 (2015). doi:10.1016/j.cois.2015.10.009

21. Dogantzis, K. A. et al. Insects with similar social complexity show convergent patterns of adaptive molecular evolution. 1–8 (2018). doi:10.1038/s41598-018-28489-5

22. Taylor, B. A., Reuter, M. & Sumner, S. Patterns of reproductive differentiation and reproductive plasticity in the major evolutionary transition to superorganismality. Curr. Opin. Insect Sci. (2019).

23. Branstetter, M. et al Genomes of the Hymenoptera. Curr. Opin. Insect Sci. 25, 65–75 (2017).

24. Standage, D. S. et al. Genome, transcriptome and methylome sequencing of a primitively eusocial wasp reveal a greatly reduced DNA methylation system in a social insect. Mol. Ecol. 25, 1769–1784 (2016).

25. Toth, A. L. et al. Shared genes related to aggression, rather than chemical communication, are associated with reproductive dominance in paper wasps (Polistes metricus). BMC Genomics 15, 75 (2014).

26. Bluher, S. E., Miller, S. E. & Sheehan, M. J. Fine-scale population structure but limited genetic differentiation in a cooperatively breeding paper wasp. Genome Biol. Evol. (2020).

27. Rehan, S. M. et al. Conserved Genes Underlie Phenotypic Plasticity in an Incipiently Social Bee. Genome Biol. Evol. 10, 2749–2758 (2018).

28. Kocher, S. D. et al. The genetic basis of a social polymorphism in halictid bees. Nat. Commun. 9, (2018).

29. Shell, W. A. & Rehan, S. M. Social modularity: Conserved genes and regulatory elements underlie caste-antecedent behavioural states in an incipiently social bee. Proc. R. Soc. B Biol. Sci. (2019). doi:10.1098/rspb.2019.1815

30. Kapheim, K. M. et al. Developmental plasticity shapes social traits and selection in a facultatively eusocial bee. Proc. Natl. Acad. Sci. 202000344 (2020). doi:10.1073/pnas.2000344117

31. Taylor, D., Bentley, M. A. & Sumner, S. Social wasps as models to study the major evolutionary transition to superorganismality. Curr. Opin. Insect Sci. 28, 26–32 (2018).

32. Emms, D. M. & Kelly, S. OrthoFinder: solving fundamental biases in whole genome comparisons dramatically improves orthogroup inference accuracy. Genome Biol. (2015). doi:10.1186/s13059-015-0721-2

33. Roy-Zokan, E. M., Cunningham, C. B., Hebb, L. E., McKinney, E. C. & Moore, A. J. Vitellogenin and vitellogenin receptor gene expression is associated with male and female parenting in a subsocial insect. Proc. R. Soc. B Biol. Sci. (2015). doi:10.1098/rspb.2015.0787

34. De Smet, F. et al. Balancing false positives and false negatives for the detection of differential expression in malignancies. Br. J. Cancer (2004). doi:10.1038/sj.bjc.6602140

35. Ilicic, T. et al. Classification of low quality cells from single-cell RNA-seq data. Genome Biol. (2016). doi:10.1186/s13059-016-0888-1

36. Alquicira-Hernandez, J., Sathe, A., Ji, H. P., Nguyen, Q. & Powell, J. E. ScPred: Accurate supervised method for cell-type classification from single-cell RNA-seq data. Genome Biol. (2019). doi:10.1186/s13059-019-1862-5

37. Furey, T. S. et al. Support vector machine classification and validation of cancer tissue samples using microarray expression data. Bioinformatics (2000). doi:10.1093/bioinformatics/16.10.906

38. Taylor, B. A., Cini, A., Wyatt, C. D. R., Reuter, M. & Sumner, S. The molecular basis of socially-mediated phenotypic plasticity in a eusocial paper wasp. bioRxiv 2020.07.15.203943 (2020). doi:10.1101/2020.07.15.203943

39. Hoffmann, K., Gowin, J., Hartfelder, K. & Korb, J. The Scent of Royalty: A P450 Gene Signals Reproductive Status in a Social Insect. Mol. Biol. Evol. 31, 2689–2696 (2014).

40. Calkins, T. L. et al. Brain gene expression analyses in virgin and mated queens of fire ants reveal mating-independent and socially regulated changes. Ecol. Evol. (2018). doi:10.1002/ece3.3976

41. Iovinella, I. et al. Antennal protein profile in honeybees: Caste and task matter more than age. Front. Physiol. (2018). doi:10.3389/fphys.2018.00748

42. Glastad, K. M. et al. Epigenetic Regulator CoREST Controls Social Behavior in Ants. Mol. Cell(2020). doi:10.1016/j.molcel.2019.10.012

43. Turrel, O., Goguel, V. & Preat, T. Drosophila neprilysin 1 rescues memory deficits caused by amyloid-β peptide. J. Neurosci. (2017). doi:10.1523/JNEUROSCI.1634-17.2017

44. Yanay, C., Morpurgo, N. & Linial, M. Evolution of insect proteomes: Insights into synapse organization and synaptic vesicle life cycle. Genome Biol. (2008). doi:10.1186/gb-2008-9-2-r27

45. Helanterä, H. An organismal perpective on the evolution of insect societies. Front. Ecol. Evol. 4, 1–12 (2016).

46. Noll, F. B., Zucchi, R. & Simões, D. Morphological caste differences in the neotropical swarm-founding polistinae wasps: Agelaia m. a. multipicta and a. p. pallipes (hymenoptera vespidae). Ethol. Ecol. Evol. (1997). doi:10.1080/08927014.1997.9522878

47. Kohannim, O. et al. Boosting power for clinical trials using classifiers based on multiple biomarkers. Neurobiol. Aging (2010). doi:10.1016/j.neurobiolaging.2010.04.022

48. Montagna, T. S. & Antonialli, W. F. Morphological differences between reproductive and non-reproductive females in the social wasp Mischocyttarus consimilis Zikán (Hymenoptera: Vespidae). Sociobiology 63, 693–698 (2016).

49. Jeanne, R. L. Social Biology of the neotropical wasp, Mischocyttarus drewseni. Bull. Museum Comp. Zool. 144, 63–150 (1972).

50. Murakami, A. S. N., Shima, S. N. & Desuó, I. C. More than one inseminated female in colonies of the independent-founding wasp Mischocyttarus cassununga von Ihering (Hymenoptera, Vespidae). Rev. Bras. Entomol. 53, 653–662 (2009).

51. Reeve, H. K. Polistes. in *The Social Biology of Wasps* (ed. Ross KG, M. R.) 99–148 (Cornell University Press, 1991).

52. Richards, O. W. The social wasps of the Americas. (1978).

53. Noll, F. B., Wenzel, J. W. & Zucchi, R. Evolution of Caste in Neotropical Swarm-Founding Wasps (Hymenoptera: Evolution of Caste in Neotropical Swarm-Founding Wasps (Hymenoptera: Vespidae; Epiponini). Am. Museum Novit. 3467, 1–24 (2004).

54. Gelin, L. F. F. et al. Morphological Caste Studies In The Neotropical Swarm-Founding Polistinae Wasp Angiopolybia pallens (Lepeletier) (Hymenoptera : Vespidae). Neotrop. Entomol. 37, 691–701 (2008).

55. West-Eberhard, M. J. Temporary Queens in Metapolybia Wasps : Nonreproductive Helpers Without Altruism? Science (80-.). 200, 441–443 (1978).

56. Sakagami, S. F., Zucchi, R., Yamane, S., Noll, F. B. & Camargo, J. M. P. Morphological caste differences in Agelaia vicina, the neotropical swarm-founding polistine wasp with the largest colony size among social wasps (Hymenoptera: Vespidae). Sociobiology 28, 207–223 (1996).

57. V., B. M., Noll, F. B. & Zucchi, R. Morphological Caste Differences and Non-Sterility of Workers in Brachygastra augusti (Hymenoptera, Vespidae, Epiponini), a Neotropical Swarm-Founding Wasp Author (s): J. New York Entomol. Soc. 111, 242–252 (2003).

58. Loope, K. J., Chien, C. & Juhl, M. Colony size is linked to paternity frequency and paternity skew in yellowjacket wasps and hornets. BMC Evol. Biol. 14, 1–12 (2014).

59. Hastings, M. D., Queller, D. C., Eischen, F. & Strassmann, J. E. Kin selection, relatedness, and worker control of reproduction in a large-colony epiponine wasp, Brachygastra mellifica. Behav. Ecol. 9, 573–581 (1998).

60. Gobbi, N., Noll, F. B. & Penna, M. A. H. ‘Winter’ aggregations, colony cycle, and seasonal phenotypic change in the paper wasp Polistes versicolor in subtropical Brazil. Naturwissenschaften (2006). doi:10.1007/s00114-006-0140-z

61. Kronauer, D. J. & Libbrecht, R. Back to the roots: the importance of using simple insect societies to understand the molecular basis of complex social life. Curr. Opin. Insect Sci. 28, 33–39 (2018).

62. Bolger, A. M., Lohse, M. & Usadel, B. Trimmomatic: A flexible trimmer for Illumina sequence data. Bioinformatics (2014). doi:10.1093/bioinformatics/btu170

63. Haas, B. J. et al. De novo transcript sequence reconstruction from RNA-seq using the Trinity platform for reference generation and analysis. Nat. Protoc. 8, 1494 (2013).

64. DI Tommaso, P. et al. Nextflow enables reproducible computational workflows. Nature Biotechnology (2017). doi:10.1038/nbt.3820

65. Buchfink, B., Xie, C. & Huson, D. H. Fast and sensitive protein alignment using DIAMOND. Nat. Methods 12, 59 (2014).

66. Li, B. & Dewey, C. N. RSEM: accurate transcript quantification from RNA-Seq data with or without a reference genome. BMC Bioinformatics 12, 323 (2011).

67. Langmead, B. & Salzberg, S. L. Fast gapped-read alignment with Bowtie 2. Nat. Methods (2012). doi:10.1038/nmeth.1923

68. Robinson, M. D., McCarthy, D. J. & Smyth, G. K. edgeR: a Bioconductor package for differential expression analysis of digital gene expression data. Bioinformatics 26, 139–140 (2010).

69. Emms, D. M. & Kelly, S. OrthoFinder2: fast and accurate phylogenomic orthology analysis from gene sequences. bioRxiv (2018). doi:10.1101/466201

70. Price, M. N., Dehal, P. S. & Arkin, A. P. FastTree 2 - Approximately maximum-likelihood trees for large alignments. PLoS One (2010). doi:10.1371/journal.pone.0009490

71. Karagiannopoulos, M., Anyfantis, D., Kotsiantis, S. B. & Pintelas, P. E. Feature Selection for Regression Problems. 8th Hell. Eur. Res. Comput. Math. its Appl. HERCMA 2007(2007). doi:10.1109/ICDM.2014.63

72. Dimitriadou, E., Hornik, K., Leisch, F., Meyer, D. & Weingessel, A. e1071: Misc Functions of the Department of Statistics (e1071), TU Wien. R package version (2011).

73. Alexa, A. & Rahnenführer, J. Gene set enrichment analysis with topGO. Bioconductor Improv. (2007).

74. Subramanian, A. et al. Gene set enrichment analysis: A knowledge-based approach for interpreting genome-wide expression profiles. Proc. Natl. Acad. Sci. U. S. A. (2005). doi:10.1073/pnas.0506580102

75. Götz, S. et al. High-throughput functional annotation and data mining with the Blast2GO suite. Nucleic Acids Res. (2008). doi:10.1093/nar/gkn176

76. Altschul, S. F., Gish, W., Miller, W., Myers, E. W. & Lipman, D. J. Basic local alignment search tool. J. Mol. Biol. (1990). doi:10.1016/S0022-2836(05)80360-2

77. Piekarski, P. K., Carpenter, J. M., Lemmon, A. R., Lemmon, E. M. & Sharanowski, B. J. Phylogenomic evidence overturns current conceptions of social evolution in wasps (vespidae). Mol. Biol. Evol. 35, 2097–2109 (2018).

78. O’Donnell, S. Reproductive caste determination in eusocial wasps (Hymenoptera: Vespidae). Annu. Rev. Entomol. 43, 323–46 (1998).

79. Matsuura, M. & Yamane, S. Biology of the vespine wasps. (Springer Verlag, 1990).

80. Menezes, R. S. T., Lloyd, M. W. & Brady, S. G. Phylogenomics indicates Amazonia as the major source of Neotropical swarm-founding social wasp diversity. Proc. R. Soc. B (2020).

