## Supplementary Table Legends for "Genetic toolkit for sociality predicts castes across the spectrum of social complexity in wasps"

**Supplementary Table S1**: **Morphometrics Data.** The body parts measured were: head length (HL), head width (HW), minimum interorbital distance (MID), partial forewing length (PWL), mesoscutum length (MSL), mesoscutum width (MSW), mesosoma height (MSH), alitrunk length (AL), length of tergite I (T1L), basal width of tergite II (T2BW), apical width of tergite II (T2AW), length of tergite II (T2L), and apical height of tergite II (T2AH). In Metapolybia, we also measured pronotal width, gena width, eye width, mesoscutellar length and mana interorbital distance. For each species we then conducted ANOVA, on each colony separately and for the listed characters.

**Supplementary Table S2: RNA-seq samples used in this study.** For each species, we show the wasp and nest IDs, caste ID, Collection site, GEO ascension numbers along with the merging of data required to form a single Queen and Worker replicate per species (and names given to these merged datasets). The second tab shows Trinity assembly statistics for the nine species (default output of Trinity). See https://github.com/trinityrnaseq/trinityrnaseq/wiki/Transcriptome-Contig-Nx-and-ExN50-stats. Showing numbers of genes and transcripts per assembly. Then conventional Nx length statistics, showing lengths of transcripts at N10/20/30/40/50 (% of assembled bases). This is performed again on the longest isoform of each gene.

**Supplementary Table S3: Orthogroups.** Trinity gene IDs for each species are listed along with their grouping into orthologous gene groups (Orthogroups: OG; tabs B:J). These are followed by counts of isoforms per species per OG (Tabs K:S). Columns "T:AM" show the data after merging of three isoforms into the single most expressed isoform. These data were made using Orthofinder.

**Supplementary Table S4: Differential expression.** Each orthogroup is tested (edgeR) to determine whether the representative gene for each species is upregulated in the reproductive (UP), down-regulated (DW), not-changing significantly (NA) or without a representative gene (NA). The sum of UP, DW and total are listed. (Tab 1 [Table of Orthogroups]). This information is extracted from the individual species edgeR results, shown in the next 9 tabs: Differentially expressed genes (9 tabs names: 1 for each species). Showing edgeR results with Trinity gene names, using hard coded dispersion of 2. Columns show standard edgeR output, with log fold change values (logFC), log counts per million (logCPM), p value between queen/worker and false-discovery rate adjusted p value (fdr). Finally, we show the TopGO result for the 95 orthogroup genes differential in at least two species (using Metapolybia gene names), with a background of the genes expressed above 1 TPM (mean across all the samples) (Tab 2 [GO top genes (0.05)])

**Supplementary Table S5: Normalised training data (Used in the SVMs).** Results from the SVM model using all nine species. Tab1 shows, A: Orthogroup, B-J: Trinity gene names for each of the nine species used in the SVM model.

J:Orthogroup. L: Blast hit (Metapolybia sequence), M-AD: showing the normalised and species normalised counts, AE/AF: Linear regression p and q coefficients, and AG-AP shows default blast2go hits for Metapolybia sequences (listed in AG), showing Seq description (top blast hit), the sequence length (AI), the number of hits (limited to 5 per sequence), Blast evalue (AK), mean similarity in % (AL). Tab 2 shows the GO result (TopGO) , of the 400 significant genes across the nine species (p value <0.05). Default output of TopGO. Bonferroni used in main figures.

**Supplementary Table S6: Details of the 10 genes that overlap between the differentially expressed genes (DEGs) and SVM genes (nine species).** Details of the genes, showing the Orthogroup name, followed by Trinity gene names for each of the nine species. Then the top two blast hits, performed using blastn (online:1/11/2020) to insect database of sequences (taxid:50557). Showing Hit (Description) name, Species name (of hit), Common name, Taxid, then blast scores and query coverage (Tab1). We also show the fasta sequences for these 10 genes, using Metapolybia sequences (Tab 2).

**Supplementary Table S7: SVM results for the four subsection SVM models.** For the four subsections: complex, simple, Independent nest founding, and swarm founding training sets. Showing Orthogroup name (A), raw TPMs for each orthogroup (C-J), normalised read counts (species scaled;K-R). Finally we show the p.coefficient aand q.coeffs of the linear regression test.

**Supplementary. Table S8: TopGO statistics for the four subsection SVM models**. Default output, showing GO ID(A) and name (B), number of gene annotated with this term (C), those significant in gene set (D), those expected given the background set(E), F-N show p values with various filters or none, the fold change (O) and ontology group (either CC: Cellular component, BP: Biological process or MF: Molecular function).
